# Improved accuracy of breast volume calculation from 3D surface imaging data using statistical shape models

**DOI:** 10.1101/2020.05.13.094227

**Authors:** M. W. Göpper, J. Neubauer, Z. Kalash, G.B. Stark, F. Simunovic

**Author notes:** Corresponding author Filip Simunovic, Department of Plastic and Hand Surgery, Division of Breast Surgery, University of Freiburg, Faculty of Medicine, University of Freiburg, Germany.

## Abstract

**Background:** Three-dimensional (3D) scanning is an established method of breast volume estimation. However, this method can never be entirely precise, since the thoracic wall cannot be imaged by the surface scanner. Current methods rely on interpolation of the posterior breast border from the surrounding thoracic wall. Here, we present a novel method to calculate the posterior border and increase the accuracy of the measurement.

**Methods:** Using principal component analysis, computed tomography images were used to build a statistical shape model (SSM) of the thoracic wall. The model was fitted to 3D images and the missing thoracic wall curvature interpolated (indirect volumetry). The calculations were evaluated by ordinary least squares regression between the preoperative and postoperative volume differences and the resection weights in breast reduction surgery (N=36). Also, an SSM of the breast was developed, allowing direct volumetry. Magnetic-resonance images (MRI) and 3D scans were acquired from 5 patients in order to validate the direct 3D volumetry.

**Results:** Volumetry based on a SSM exhibited a higher determination coefficient (R^2^=0,737) than the interpolation method (R^2^=0,404). The methods were not equivalent (p=0.75), suggesting that the methods significantly differ. There was no influence of BMI on the correlation in either method. The MRI volumetry had a strong correlation with the 3D volumetry (R^2^=0,978).

**Conclusion:** The SSM-based method of posterior breast border calculation is reliable and superior to the currently used method of interpolation. It should serve as a basis of software applications aiming at calculation of breast volume from 3D surface scanning data.

## Introduction

A reliable estimate of preoperative breast size facilitates oncologic and esthetic breast procedures, breast reconstruction and reduction surgery. Breast volume calculation can assist in stratifying patients preoperatively, and in quantifying results of surgery, leading to more objective scientific reporting. Also, insurance companies rely on breast measurements to decide whether to cover the costs of treatment for breast reduction.

A number of methods are used for estimating breast volume [1-3]. Anthropomorphic methods rely on distance measurement between surface points. A Grossman-Roudner device consists of graduated discs which are converted into a cone, and the read from calibrations marked on the disc. In thermoplastic procedures a cast of a patient’s breast is produced, and the volume of the fluid which fills the cast recorded. Archimedian methods suggest that the patient submerges her breast into a basin and that the amount of displaced fluid is quantified. Breast volume has been derived from sonography, mammography, computed tomography (CT), and magnetic resonance imaging (MRI). Finally, the water displacement of a mastectomy specimen is considered to be the most exact volume measurement of a female breast. With the exception of anthropomorphic measurement and the Grossman-Roudner method, the above methods are not routinely applicable because of patient discomfort, technical complexity and costs. Anthropomorphic measurements and the Grossman-Roudner method have been shown to be acceptably accurate in a comprehensive work by Kayar et al. The main limitation of these methods is that their accuracy rapidly decreases in breasts larger than 500 ml [4, 5].

In search for a method that is convenient for the patient and the practitioner, 3D laser surface scanning has been used by several groups. 3D volumetry was shown to be accurate when compared to mastectomy specimens [6], MRI [1, 7] and thermoplastic casting and anthropomorphic measurements [8]. However, the posterior boundary of the 3D image of the breast remains an unsolved problem, since an image acquired by the surface scanner can never include the thoracic wall (**Figure 1A**). The most widely reported method of posterior wall calculation includes interpolating the boundary from the edges of the “defect” which is created when the breast is “cut out.” While feasible in normal-BMI patients, (**Figure 1B**) the method reaches its limitation in adipose patients [7], since axillary rolls and upper abdominal fat tissue preclude a reliable interpolation of the thoracic wall (**Figure 1C**).

**Figure 1.**
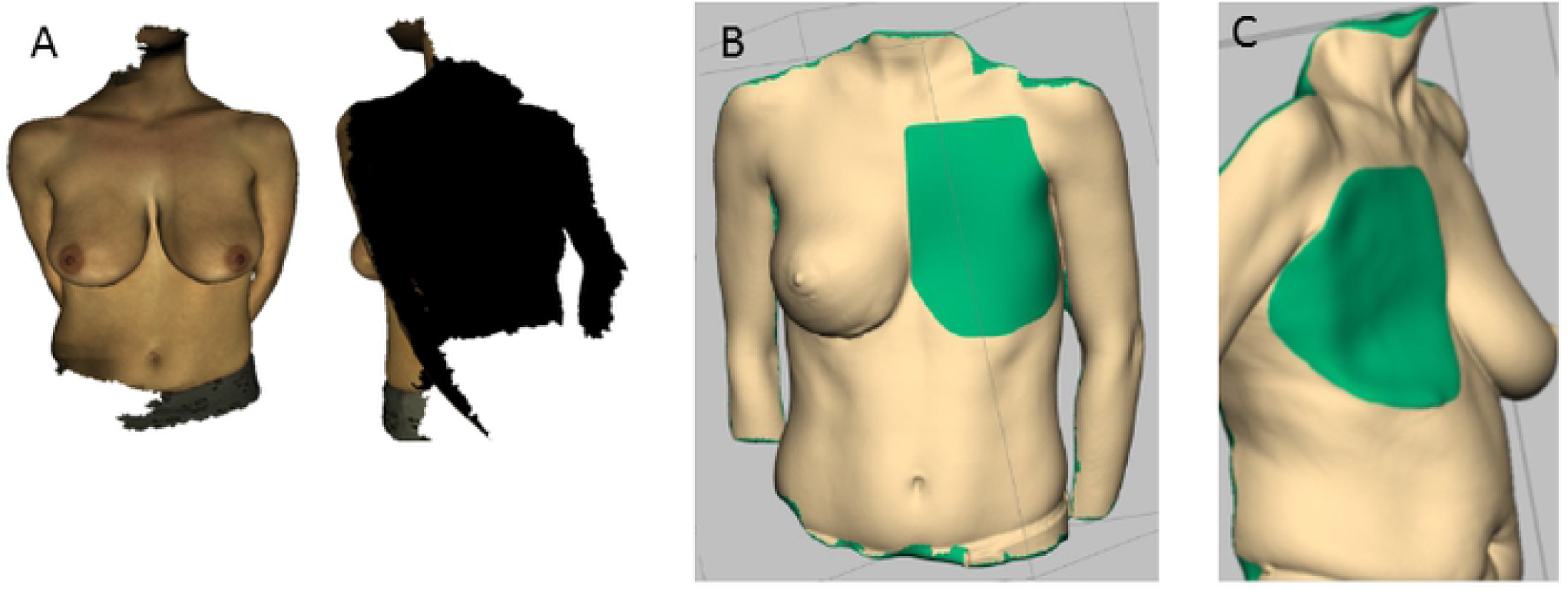
3D laser surface scanning images do not include the posterior border of the breast. (**A**). Standard methods for the estimation of the posterior border involve an attempt by a software to interpolate the missing posterior border based on the edges of the model. This method was shown to be reliable in patients with low or normal body-mass index (BMI) The person pictured has a BMI of 23 kg/m^2^ (**B**). However, since patients with higher BMI values exhibit more subcutaneous tissue at the edges of the breast, interpolation can result in a non-anatomical prediction of the thoracic wall and the accuracy of the method is reduced, as shown in this case with a BMI of 27 kg/m^2^ (**C**).

We used CT images of general population to create, by means of principal component analysis (PCA), a statistical shape model (SSM). This model was merged with 3D laser scanning data and used for interpolation of the missing thoracic wall curvature. We hypothesized that thoracic wall interpolation based on a SSM is more accurate in general, and especially in the higher-BMI population, than interpolation which relies on the edges of the breast.

## Materials and Methods

### Patients

Thirty-nine patients that underwent breast reduction surgery at our department between December 2016 and November 2017 were included in this study (**Supplementary table 1**). The mean age was 46.2 (±15.5) years and the BMI was 29.7 (±6.9) kg/m^2^. Thirty-three procedures were “inverted T” reductions with a cranially based nipple-areola-complex as described by Höhler [9], and three procedures were vertical scar procedures as described by Lejour [10]. Resection weight was measured intraoperatively. The weight (g) was converted to volume (ml) with a correction index of 1.07 for premenopausal and 1.06 for postmenopausal patients [11]. The cutoff between was set at 55 years [12]. The mean corrected resection volume was 689.8 ± 383.8 ml. The patients were scanned preoperatively and postoperatively at: 2 and 5 weeks, 6 and 12 months. Eighteen patients were seen after 6, and only 15 after 12 months. Three patients were entirely lost to follow-up or had poor quality scans. We first performed an analysis of 18 cases which were followed-up at 6 months and then on all available cases using the latest available follow-up (7 patients at 2 weeks, 7 at 5 weeks, 8 at 6 months and 14 at one year).

### 3D surface scanning

A MHT surface scanner (Artec Group, San Diego, Calif.) was used, which has a three-dimensional resolution of 0.5 mm and a three-dimensional point accuracy of up to 0.1 mm. A flashing light projects a grid pattern onto the surface, and the distortion is captured by three cameras from different. The data is imported in Artec Studio Professional (Version 12, Artec 3D, Luxemburg, **Figure 3A**), and the area of the breast removed from the scan (**Figure 3B and C**), based on defined anatomical landmarks: the medial border is the midsternal line, the caudal border is one centimeter below the submammary fold, laterally it incorporates the anterior axillary fold, and cranially the border is a straight line connecting the jugulum with the anterior axillary fold. These borders were also used in previous publications [7].

**Figure 2.**
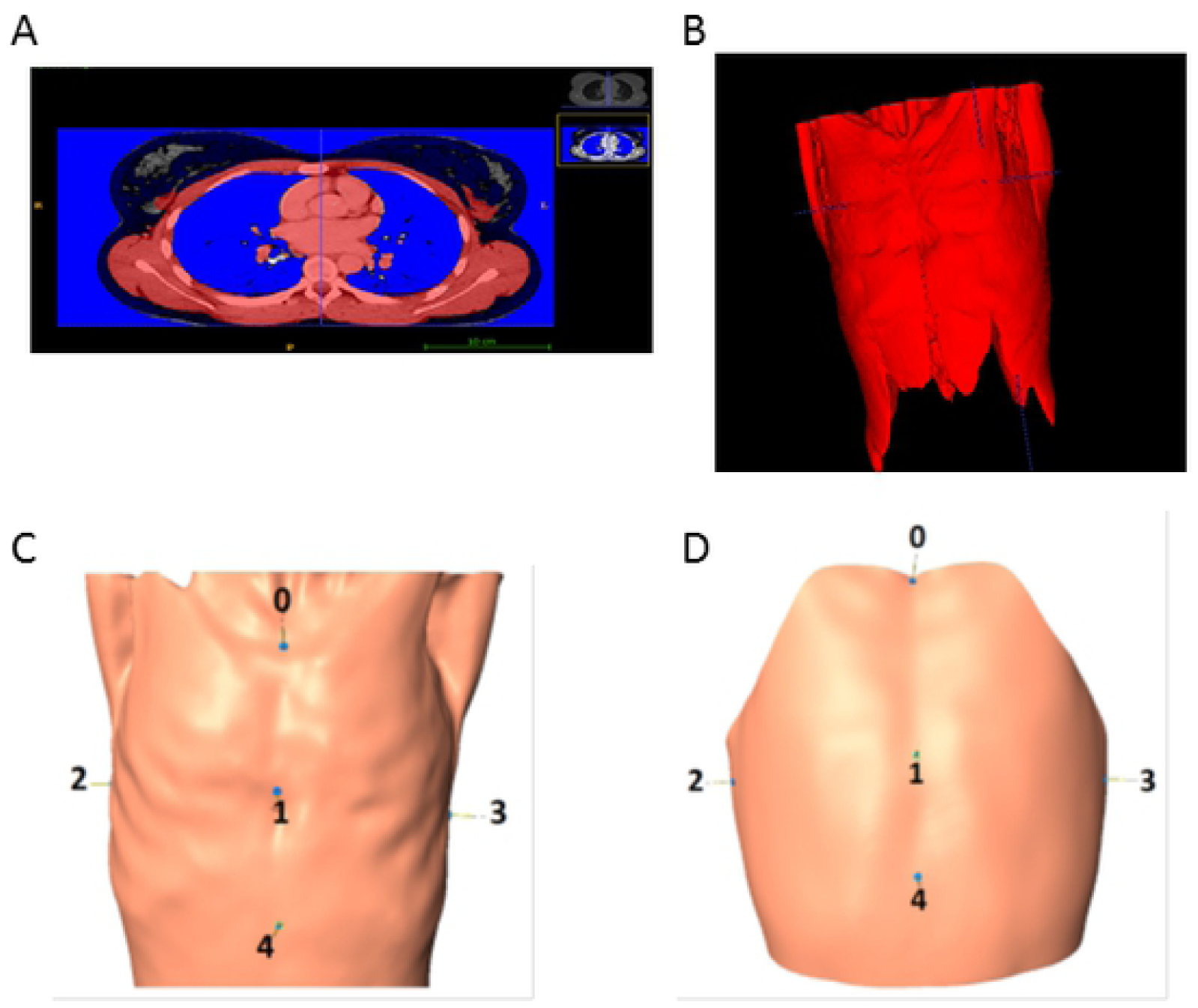
Computed tomography (CT) image data is processed in order to incorporate it into a statistical shape model (SSM). The CTs are from 72 female patients which received CTs for other reasons and were not suffering from thoracic pathologies. Image segmentation (**A**) was performed to create images with closed surfaces (**B**), and this data was used to build a statistical form model. Anterior and posterior axillary folds were removed, and 5 standardised landmark points were placed on the model (0 – jugulum, 1 – a point between the caudal and the middle third of the sternum, 2 – an intersection of the fourth rib an the posterior axillary fold on the left, and 3 - right sides, 5 – a point 10 centimeters below the xyphoid, **C** and **D**).

**Figure 3.**
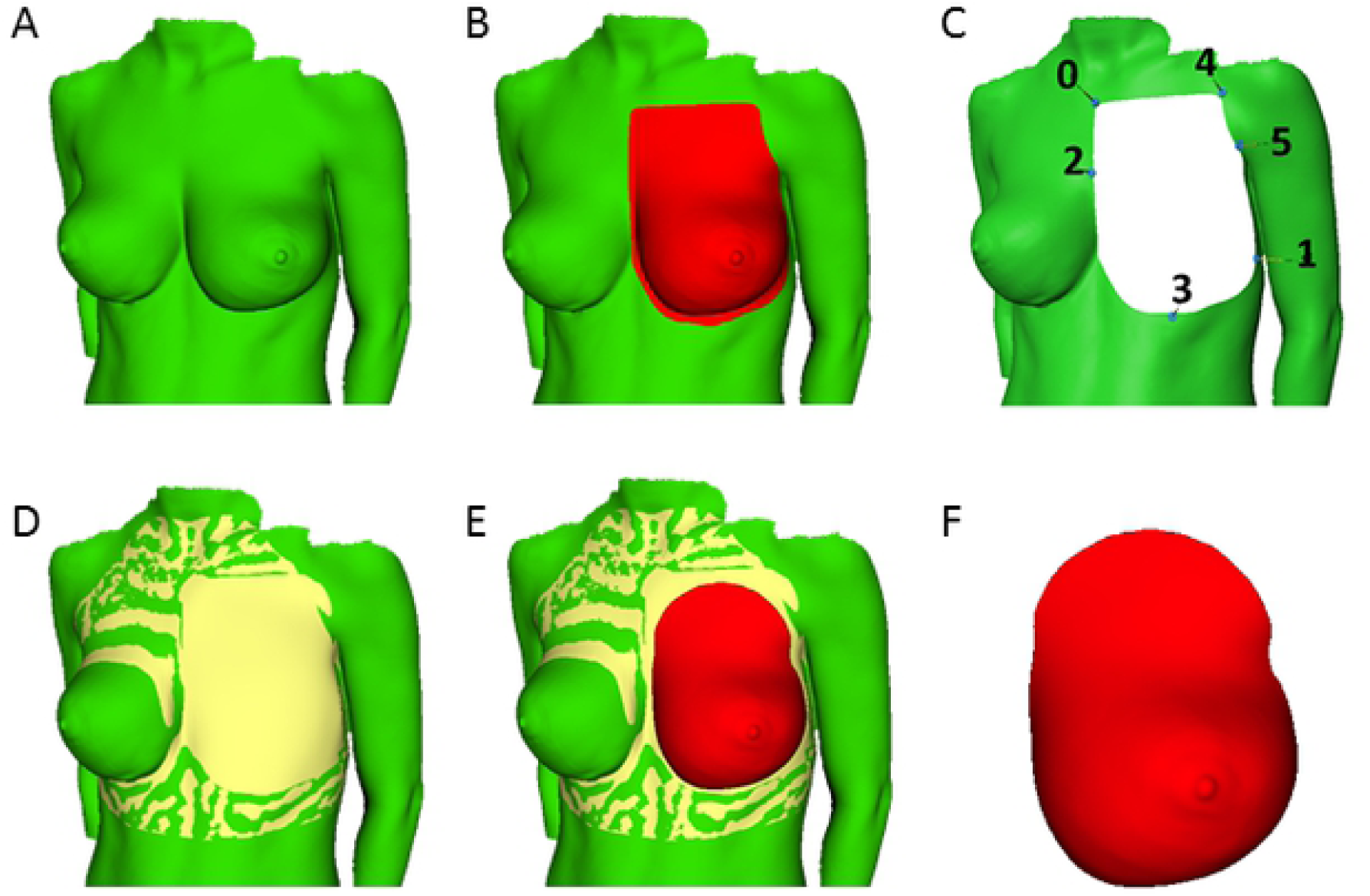
The statistical shape models (SSM) derived from CT scans and 3D surface scanning data are integrated. Six landmarks are manually placed on the model to allow for initial rotation (**A to C**). To reduce the interobserver error and semi-automatically identify the configuration with the closest point distribution, an iterative closest point algorithm (ICP) is run after rotation. Afterwards, elastic registration is performed to achieve a better fit on the cutting edges of the target mesh. After fusion of the thoracic wall model and the non-preprocessed surface scan (**D and E**), a 3D mesh of the breast is acquired that can be used for volume calculation (**F**).

### Principal component analysis was used to build a statistical form model

SSM was built from segmented 3D data of CT images of 72 patients which underwent whole-body CT scans at the emergency center of the Freiburg University Medical Center for polytrauma diagnostic workup. We selected female patients which had no thoracic pathology and divided these in six age groups: <20 years (N=11), 20-29 years (N=10), 30-39 years (N=13), 40-49 years (N=12) and >60 years (N=16). The segmentation (**Figure 2 A and B**) was performed with the Insight Segmentation and Registration Toolkit [13] and its active contour algorithm [14]. Meshes from the segmented CT data were registered by a freeform deformation based on 5 landmarks. For model building we used the R – Packages Rvcg, Morpho, mesheR and RvtkStatismo [15-20].

Six landmarks are placed on the model to allow for initial rotation (**Figures 3 A to C**). An iterative closest point algorithm [21] is run to reduce the interobserver error, and 20 fitting steps are performed with elastic registration [22] to achieve a better fit. After fusion of the thoracic wall model and the surface scan (**Figures 3D and E**), a 3D mesh of the breast can be used for volume calculation (**Figure 3F**).

The interpolation of the posterior border from the edges of the thorax was performed as previously described [7] using Geomagic Freeform Plus (Raindrop Geomagic, Inc, NC)

All 3D scans from the study (N=130) were used to create a SSM of the breast, which already includes the posterior breast border. Instead of closing the posterior breast border on the model of the thorax, the posterior breast border is closed on the model of the breast. The SSM of the breast simplifies the user experience (**Figure 5A-B**).

**Figure 4.**
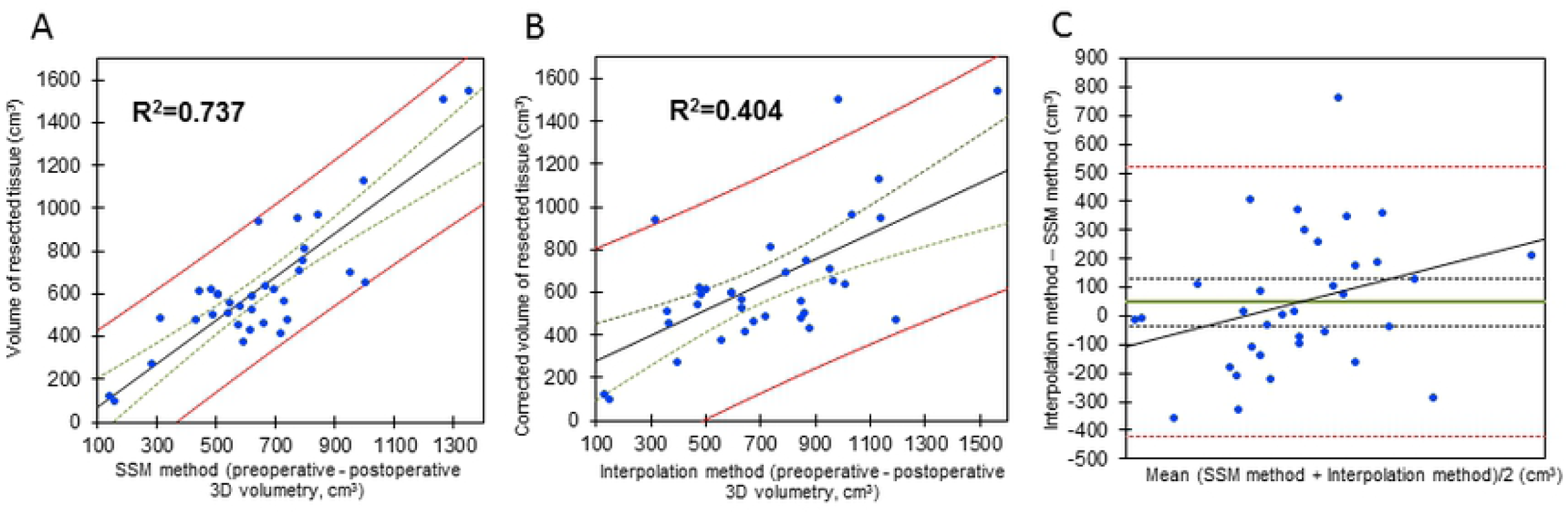
Ordinary least square (OLS) regression analysis showing that the statistical shape model (SSM) - based estimation of breast volume has a stronger correlation with the resection weight than the interpolation-based method. The correlation of the resection weight in breast reduction surgery and the difference between the preoperative and postoperative scans (N=18 patients, 36 breasts) was found to be higher when the posterior border was interpolated with the SSM-based method (**A**, R^2^=0.737) as compared to interpolation which relied on edges of the breast (**B**, R^2^=0.404). This calculation includes the patients which had followup scans at 6 months. **C** shows a bland-altman plot comparing the SSM-based and the interpolation method. With a total p-value of 0.75 in TOST (two-one-sided-test), the equivalence between the methods is rejected, signifying a difference between methods. The green lines signify 25% and the red lines 95% confidence intervals.

**Figure 5.**
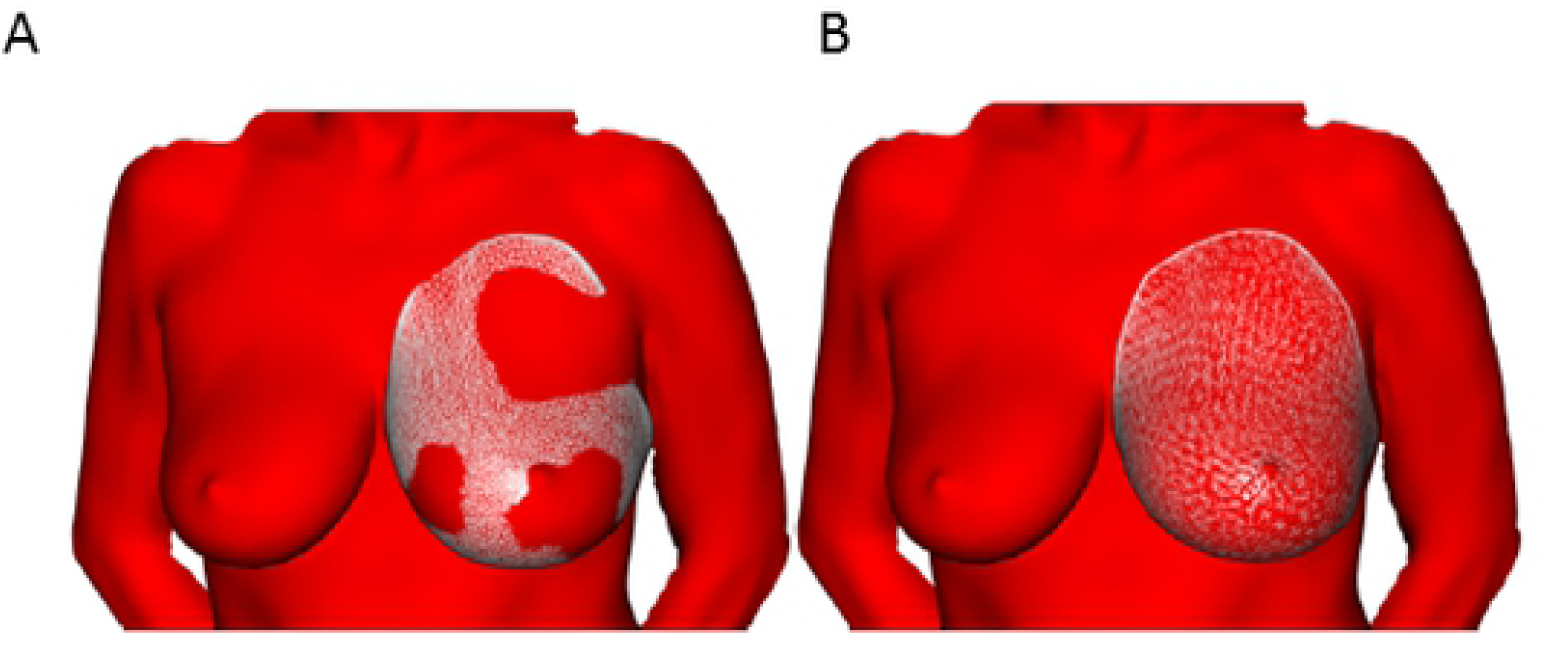
A statistical shape model (SSM) of the breast allows direct volumetry and improves the applicability of the method. SSM of the breast includes the posterior breast border. In other words, instead of closing the posterior breast border on the model of the thorax, the posterior breast border is closed on the model of the breast. The SSM enables the researcher, after manually selecting the predefined landmarks on the 3D scan, to fit the model onto the actual 3D scan (**Figure 5A**), with subsequential elastic registration to achieve a better fit (**Figure 5B**). This makes the process significantly more user-friendly. See also **supplementary figure 2**.

### SSM-based direct volumetry: user workflow

Scanned images are first scaled in MeshLab [23] or Artec Studio Professional (Version 12, Artec 3D, Luxemburg). These are further processed in Landmark software (Institute for Data Analysis and Visualisation, Davis, CA), where points are manually placed on the breast of interest (**supplementary figure 3**). These points are the basis for fitting the SSM of the breast onto the 3D scan of interest. The volume is then calculated in R-Studio (RStudio, Inc; Boston, MA).

### Magnetic resonance imaging and breast volumetry

Five patients were examined in a 3T scanner (Magnetom VIDA, Siemens Healthcare GmbH, Forchheim, Germany), placed in a prone position and the breasts positioned in bilateral 18-channel breast coils. A transverse T1 DIXON sequence was performed with a slice thickness of 2 mm, field of view of 340 × 340 mm, matrix of 576 × 461, TE of 2.46 and TR of 5.96. For processing we used 3D slicer to perform an Otsu-threshold segmentation in the gray scale range from 200 to 700 [24-26], which was smoothed with a 10 mm kernel (**Supplementary figure 4 A and B**).

### Statistical analysis and ethical approval

Fisher’s F test and the Kolmogorov-Smirnov tests were performed to compare the variances between the left and right sides. The tests showed no differences in distributions, and both sides were combined in further analysis. Ordinary least square (OLS) analysis was used to calculate the regressions. The equivalence between the two methods was tested by a Bland - Altmann plot [27] and a two-one-sided-test (TOST) for two independent samples. The calculations were performed using XLSTAST (Addinsoft 2020) and Stata/IC (version 12.1., StataCorp LP, 4905 Lakeway Drive, College Station TX 77845, USA). Results were considered significant if p < 0.05.

All parts of the study have been approved by the Ethics Committee of the Medical Faculty, Freiburg University, and all patients have provided informed consent for participation in the study.

## Results

### The SSM-based posterior border estimation of breast volume has a stronger correlation with the resection weight than the interpolation-based method

An OLS analysis on 18 patients with a 6 month follow-up showed an R^2^ of 0.737 (p<0.01) for the SSM method (**Supplementary table 2**) and an R^2^ of 0.405 (p<0.01) for the interpolation method (**Supplementary table 3** and **Figure 4 A** and **B**). The equivalence between the two methods was rejected, (p= 0.75), suggesting that the two methods significantly differ (**Figure 4C**).

When repeating the analysis on 36 patients which had different follow-up time points, again a higher coefficient of determination (R^2^ = 0.776, p<0.01) for the SSM-based method of posterior border evaluation was found than for the interpolation method (R^2^ = 0.594, p<0.01; **supplementary figure 1 and supplementary tables 4** and **5**). The equivalence between the two methods was also rejected with a p-value of 0.824.

### The body-mass-index does not interact with the correlation between the resection weight and the estimated breast volume

While the BMI significantly correlated with the resection weights in all groups, there was no significant interaction of BMI on the correlation between resection weight and volumetric volume resection. The interaction coefficients were 0.007 for the SSM and 0.008 for the interpolation method in the first patient group (6 month follow up), and - 0.003 in the PCA and 0.003 in the interpolation method in the second patient group (latest available follow up, **supplementary tables 2-5**). All interactions had a p>0.05.

### Direct breast volume estimation based on a statistical shape model with posterior border and validation by magnetic resonance imaging

The 3D scans that were acquired in the course of the study were used to create a statistical shape model of the breast, meaning that the model already includes the posterior border, allowing direct volumetry of the breast (**Figure 5**). We compared estimated volumes for five 3D surface scans that were not part of the model building in the previous described parts of this paper. We also have MRI-volumetric data of these patients so that we were able to calculate a logistic regression. An OLS analysis showed an R^2^ of 0.953 (p<0.01) for the SSM-based direct volumetry (**Figure 6 A**) and an R^2^ of 0.978 (p<0.01) for the SSM-based thoracic wall estimation method formerly used for indirect volumetry (**Figure 6 B**). We conclude that direct SSM-based volumetry leads to similar results than SMM-based estimation of the posterior border of the breast but is much more user-friendly.

**Figure 6.**
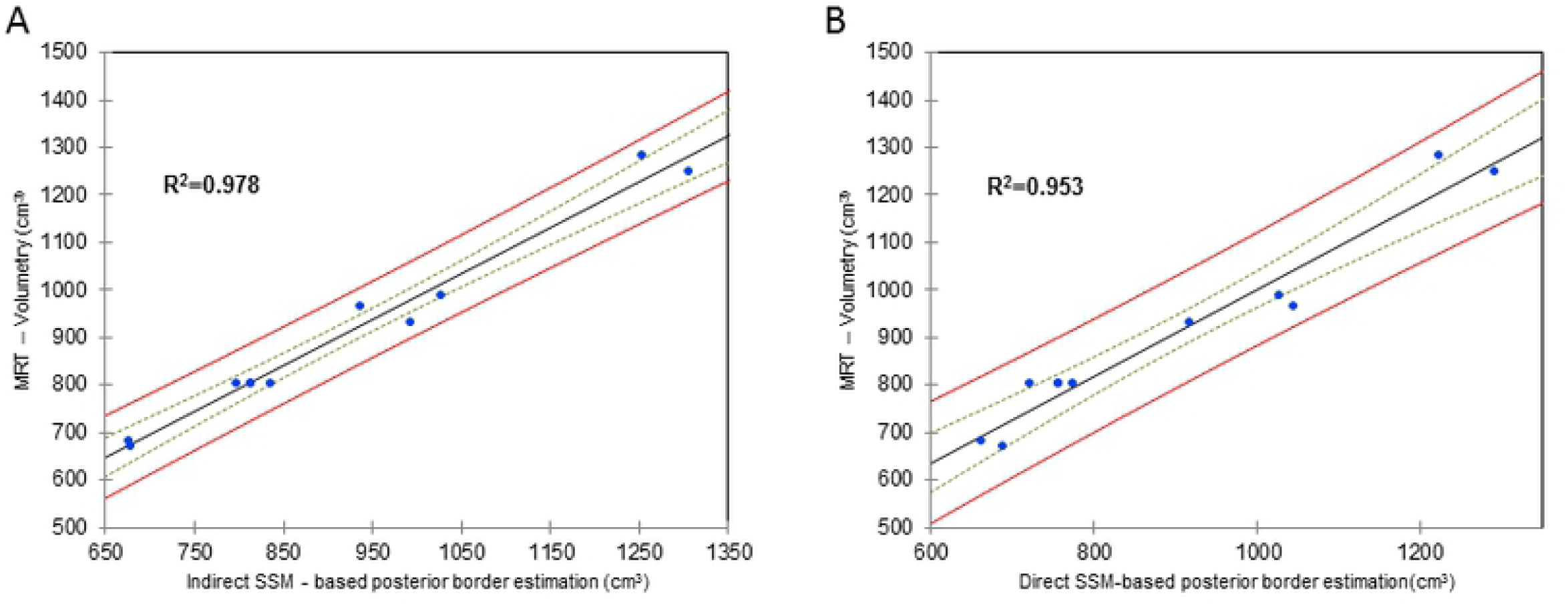
3D volumetry is validated by magnetic resonance imaging (MRI). In **A**, MRI volumetry is plotted against the SSM - based posterior border estimation methodshowing a very high correlation of R^2^=0.978. In (**B**) theaccuracy of the direct SSM-based volumetry of the breast with integrated posterior border was validated by MRI volumetry (**B**) where a correlation of R^2^=0.953 has been achieved. The green lines signify 25% and the red lines 95% confidence intervals.

## Discussion

### Breast volumetry is of key importance in breast surgery

Breast surgery is marred by lack of objectifiable parameters, and surgical techniques are difficult to describe and teach [28]. The lack of objective criteria also hampers scientific progress. For example, the “Breast cancer conservative treatment score” (BCCT.core) is a useful tool for calculating a symmetry index between the breasts [29]. The Telemark Breast Score also aims at standardising photographic material to deliver a score of symmetry, volume and shape [30]. The ideal software should however contain (largely) automatic shape, symmetry and volume measurement. The plethora of methods testifies that especially the latter is of great interest, since volume quantification can facilitate stratification of patients and operative planing. With more accessible 3D surface scanning devices, this technology has been used by several groups for breast volume quantification. 3D volumetry was correlated to MRI [1, 7, 31, 32], thermoplastic casting and anthropometric measurements [8], and mastectomy specimens [6]. 3D imaging has also been used to investigate postoperative courses. Eder et al. showed that posoperative changes after breast reduction continue for 3-6 months [33], and after breast augmentation for 3 months [34], while another group suggested 6 months [35]. Others have focused on symmetry, finding a high grade of breast asymmetry in general population [36], and in 92 out of 100 patients after augmentation [37].

A review of literature concluded that 3D imaging and MRI are the most reliable methods of breast volumetry. The authors conclude that variance between the studies is due to definitions of the borders, and especially of the posterior breast border [38]. The standard method of posterior border estimation is interpolation from the thoracic borders (**Figure 3C**) using dedicated software (e.g. Raindrop Geomagic, Inc, NC). The studies were done on normal BMI patients (e.g. 20 kg/m^2^) and the colleagues conclude that the method might be less reliable in overweight patients [7]. We postulated that by creating a statistical form model based on CT images of the thorax a more anatomical interpolation of the thoracic wall would be achieved.

### SSM predict datapoints not contained in the original dataset

PCA is a statistical procedure used for creating predictive models, such as SSM. 3D meshes (segmented CT data and surface scans) are described as a set of coordinates in three dimensions. Sets of these coordinates are statistically processed and analyzed using PCA [39]. If normal distribution is given, transposition of the covariance matrix leads to diagonal size sorted vectors, with each vector describing one principal component. We hypothesized that the precision of the volumetric estimation would increase when using an SSM-based posterior border prediction.

### SSM-based posterior border calcutation is reliable and BMI-independent

Methods of posterior border interpolation (SSM-based vs. interpolation) were evaluated by correlating the volume differences preoperatively and 6 months postoperatively. The 6 month timepoint was chosen because the evolution of postoperative changes (edema, seroma, haematoma) is usually resolved and the final breast shape reached [33]. This method of validation can be confounded by many factors: menstrual cycle, changes in weight, postoperative swelling, edema and hematoma. However, this method corresponds to the most exact method of volume determination (water displacement of the mastectomy sample) and is superior to any other non-radiological method of breast volumetry. In the second part of the study, SSM-based direct volumetry was validated by MRI-volumetry.

There was no influence of BMI on the regression analysis in either of the methods. The analysis was performed on a population which was overweigt (29.7 kg/m^2^) and had a large enough standard deviation (±6.9 kg/m^2^) to sufficiently reflect a variety of habitus.

### Conclusions

The Artec MHT surface scanner is a large device that requires a power source and that needs to be connected to a computer. This setup is inconvenient, so that we currently use the Crisalix 3D surface Imager Camera (Crisalix S.A., Lausanne, Switzerland) mounted on an iPad tablet (Apple Inc., Cupertino, CA). The Artec scanner delivers more information to the 3D dataset; however, this is not needed for the calculation of the breast volume. Also, the cheaper price and increased convenience of use of the Crisalix system prevail. The Crisalix software (Crisalix S.A., Lausanne, Switzerland) also calculates breast volume, in adittion to its main function to simulate postoperative outcomes. Since we do not consider such simulations reliable or useful for patient counceling, and due to an exorbitant price tag, we have no experience with the software. A recent study has suggested that 3D volume estimation with the Crisalix scanner and Crisalix software is accurate in breasts of <600 cm^3^ when compared to surgeon estimates, anthropometric measurements, and mastectomy speciments [40]. More experience is needed to prove the accuracy of the Crisalix software in breast volume calculation. At the moment, 3D laser scanning volumetry is routinely used at our department for preoperative volume determination in breast reconstruction and reduction cases. The current workflow to generate a volume estimate from a 3D scan involves three steps in three separate software applications (see “SSM-based direct volumetry: workflow” in the methods section) and is thus not easily transferable to other practitioners. Another issue is that especially the second step, which involves manually selecting the points of measurement on the borders of the breast (**Supplementary figure 3**), is susceptible to inter-user variability [6, 41]. The effect of interindividual variance is reduced by elastic registration, which compensates for deviations between points (**Figure 5**). An automated measurement of breast volume has been attempted from MRI images and 3D scans [42].

We present the most precise 3D volumetric method to date. Every measurement, however, has a range of precision; and advances in science are not made by continuously trying to reduce this range to nothing [43]. Even if we knew the absolute volume of the breast, this knowledge would only marginaly improve our practice, as it is rather the relation of the volumes (between the breasts, or between preoperative and postoperative measurements) which is of interest to us. Further advances in design of tools for objective quantification of the female breast will not arise from further refinement of volume measurement, but rather in integrating the methods of volume determination based on 3D scanning with mesures of breast symmetry and shape in an acessible and user-friendly software environment.

## Competing interests

The authors have no conflict of interest to report

### Acknowledgments

The authors wish to acknowledge Dr. Pinar Simunovic for her help with data analysis.

## Figure Legends

**Supplementary figure 1.**
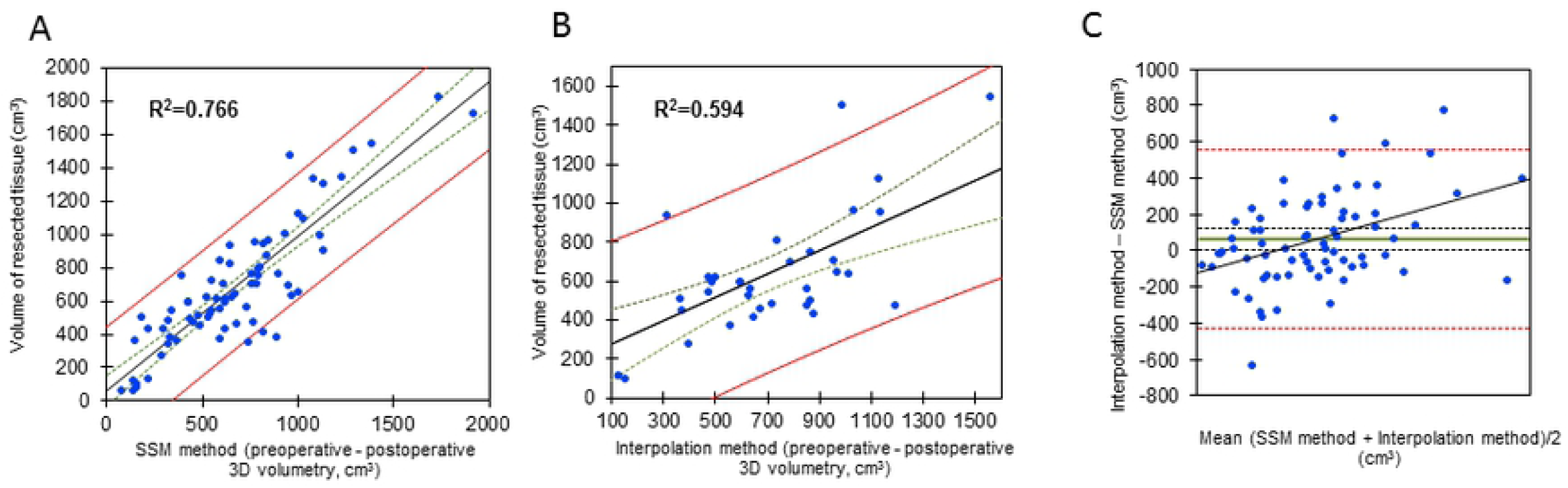
Ordinary least square (OLS) regression analysis showing that the statistical shape model (SSM)-based estimation of breast volume has a stronger correlation with the resection weight than the interpolation-based method. The correlation of the resection weight in breast reduction surgery and the difference between the preoperative and postoperative scans (N=36 patients, 72 breasts) was found to be higher when the posterior border was interpolated with the SSM-method (**A**, R^2^=0.766) as compared to interpolation which relied on edges of the breast (**B**, R^2^=0.594). In this analysis, the latest possible followup timepoint was selected, which means that the followup timepoints are heterogenious. abland-altman plot is drawn comparing the SSM-based and the interpolation method (**C**). .The plot shows some values exceeding the 95% confidence interval. Furthermore, the values seem to cluster at low values and again some kind of logistic regression is noticeable so that we state difference between the comparised methods. When comparing the two methods, we see that with a total p-value of 0.824 in TOST, the equivalence between the methods is rejected, suggesting a significant difference between the SSM-based and the interpolation method. The green lines signify 25% and the red lines 95% confidence intervals.

**Supplementary figure 2. Creation of a statistical shape model (SSM) of the breast**. While the first part of the study was focused on creating an SSM of the thoracic wall (**Figure 2**), we aimed at designing an SSM of the breast for purpose of direct breast volumetry. This model already contains the SSM-based prediction of the posterior breast border and greatly simplifies the workflow, since it becomes sufficient to mark the points on the scan of breast (**supplementary figure 3**), instead of having to mark the whole thorax. Samples for building up the model are the estimated breasts of patients participating in this study. Furthermore, all pre- and postoperatively measured patients were included in the model, so that more data than just indirect volumetry were available, thus increasing the statistical coverage of the SSM. Overall, there are 130 samples per page.The above images show the placement of the points for the registration of the samples by a free form deformation based on 6 individually set landmarks for initial rotation. Model building was performed analogous to the SSM of the thoracic wall: Regularization was achieved by a Gaussian Process model that was created from a generic breast template. To allow for shapes outside the model space, a weighted average between the free form deformation and the model was calculated, with the model weights decreasing by each iteration (to 90% of the weight of the previous iteration). The lower two images are illustrating the the elastic registration process. Again we get shapes where each point is in correspondence to each other and we are able to define the covariance matrix which is essential generating a SSM. The procedure has to be done seperately for the left and the right breast.

**Supplementary figure 3.** Points are manually placed on the breast of interest. These are the basis for the merge of the 3D scan of the patient with the statistic form model, leading to a 3D breast image with closed posterior border. Manual placement of the points renders this step voulnerable to interuser variability. To reduce the interobserver error and semi-automatic identify the configuration with the closest point distribution, an iterative closest point algorithm (ICP) is run. Afterwards 20 fitting steps are performed by reducing the MSE (mean squared error) from model to target with subsequential elastic registration to achieve a better fit. Anatomical definitions of the points are: 0 – nipple, 1 – cranial breast border (curved extension of the preaxillary fullness), 2 – medial breast border, 3 – caudal border (in the submammary fold), 4 – lateral border of the breast (usually the anterior axillary line), 5 – point halfway between 0 and 1, 6 - point halfway between 0 and 2, 7 - point halfway between 0 and 3, 8 point halfway between 0 and 4, 9 – mediocranial border (point halfway on the curved line between 1 and 2), 10 – laterocranial border (breast border in the anterior axillary line), 11 – laterocaudal border (point halfway between 3 and 4).

**Supplementary figure 4. MRI-segmentation for MRI-volumetry** To process the MRI data we used 3D slicer to perform a Otsu-threshold segmentation in the gray scale range from 200 (lower threshold) to 700 or maximum value if below 700 (upper threshold) (**A**). The resulting segmentation was then smoothed using in 3D slicer implemented Gaussian filters with a 10 mm kernel in the threshold range from 50 to 700. Subsequently and the edges were dilated with a 10 mm kernel in the threshold range from 50 to 700 too. The result is a manually adjusted segmentation with a boundary to the chest posterior wall at skin level (**B**).

